# The Effect of Lipid Composition on the Dynamics of Tau Fibrils

**DOI:** 10.1101/2022.02.16.480652

**Authors:** Unmesh D. Chowdhury, Arnav Paul, B. L. Bhargava

**Affiliations:** School of Chemical Sciences, National Institute of Science Education & Research-Bhubaneswar, an OCC of Homi Bhabha National Institute, P.O.Jatni, Khurda, Odisha 752050,India

## Abstract

Knowledge of the interaction of the tau fibrils with the cell membrane is critical for the understanding of the underlying tauopathy pathogenesis. Lipid composition is found to effect the conformational ensemble of the tau fibrils. Using coarse grained and all-atom molecular dynamics simulations we have shown the effect of the lipid composition in modulating the tau structure and dynamics. Molecular dynamics simulations show that tau proteins interact differentially with the zwitterionic compared to the charged lipid membranes. The negatively charged POPG lipid membranes increase the binding affinity of the tau fibrils. The addition of cholesterol is also found to modify the tau binding to the membrane.

## 1 Introduction

Tauopathies comprises of a set of central nervous system diseases such as Alzheimer’s disease (AD), frontotemporal lobar degeneration, Pick’s disease, and progressive supranuclear palsy, characterized by abnormal aggregation of the microtubule-associated protein tau (MAPT).^1^ Tau protein polymerizes tubulin to form microtubules (MT) and provides axonal support to MTs. Tau is an intrinsically disordered protein (IDP) that is soluble in normal conditions but can form insoluble aggregates. In a healthy brain tau proteins undergo phosphorylation which is 2-3 moles of phosphate per mole of tau protein. Hyper-phosphorylation in tau is when the phosphate mole ratio is almost 3 fold more than a normal brain tau. ^2^ Hyperphosphorylation of tau makes it difficult for it to associate with MTs, and causes aggregation of tau and formation of paired helical filament (PHF) and neurofibrillary tangles (NFT).^3^

AD is characterized by gradual accumulation of amyloid plaques consisting of aggregates of A*β* peptides and aggregates of tau. Research on AD was majorly focused on A*β* peptides but due to failures of A*β* targeting treatments in clinical trials, tau proteins are getting the attention of researchers.^4^ A total of six tau protein isoforms are expressed in AD brain, ranging from 352 to 441 amino acids. The six isoforms of tau can be distinguished based on the presence or absence of inserts of 29 or 58 amino acids in the N-terminal half, and the inclusion or absence of the 31 amino acid microtubule-binding repeat in the C-terminal half. In general tau can be said to have four broad domains, the N-terminal domain, the proline rich domain, repeat domain region and the C-terminal domain. ^5^ Fetal brain has only three repeat domains (R3 tau) while the adult human brain has four repeat domains (R4 tau). The repeat domains R3 and R4 form the core structure for the paired helical filaments (PHF) and straight filaments (SF).^6^ PHFs and SFs both are composed of two protofilaments with C-shaped subunits.^6^ Cryo-electron microscopy structures of tau filaments from the individuals with AD were determined by Fitzpatrick and coworkers which serves as a good starting point for MD simulations.^7^ Studies have shown that tau interacts with plasma membrane through its amino projection domain. ^8^ Interaction of tau with plasma membrane is observed to promote tau aggregation in vitro, however, the exact mechanism is unknown.^9^ Anionic lipid membranes facilitates the fibrillation of tau and A*β* proteins.^10,11^ In the microtubule binding (MTB) region of tau, three segments (253-261, 315-323 and 346-355) have been observed to bind to lipids and take up a helical structure that facilitates protein aggregation.^12,13^ Membrane lipids like phosphatidylcholine (PC), cholesterol, and sphingolipid have been observed to be associated with the tau proteins.^14^ Membrane interactions with tau proteins show that the membranes play an important role in fibrillation and associated toxicity as reported in case of PHF6 hexapeptide.^15^

Molecular dynamics simulations help in characterizing the intrinsically disordered proteins (IDPs) which do not possess any particular three dimensional structure. ^16^ IDPs possess seemingly dynamic conformation ensemble that are usually characterised by small angle neutron/X-ray scattering (SANS/SAXS), nuclear magnetic resonance (NMR), circular dichroism, fluorescence resonance energy transfer (FRET) etc. ^17–20^ The data from the experimental techniques mentioned above are usually scarce for a complete structural and dynamical characterisation of the IDPs. ^21^ Recent development of the optimized force fields for the IDPs along with compatible water models^22–24^ have further improved their computational modeling. The interaction of lipid membrane with the amyloid fibrils are studied using both the coarse-grained and the all-atom models.^25,26^ Computer simulations have shown that in case of zwitterionic bilayers, fibril-membrane binding is dominated by coulombic interactions. ^27,28^ Jang and coworkers have shown the possible fibril conformations along the pathways for the membrane insertion. ^25^ The increased cholesterol level also hints at the improved binding of the monomeric A*β*-42 to the bilayers.^28^ In a different study using the replica exchange molecular dynamics (REMD), it was found that the cholesterol prevents the penetration of the A*β*-40 monomer into the lipid bilayer. ^29^ The hexapeptide ^306^*V QIV Y K*^311^ (PHF6) is used as a template in a number of MD simulations due to its similarity to the tau fibrils in vitro and also because its omission is found to prevent tau assembly.^30,31^ Few other MD simulations describing the conformational states of the tau helical filaments have been reported recently.^32–34^ To the best of our knowledge, there are no reports of comprehensive computational study of the entire stretch of tau proteins with various lipid compositions. In this study, we elucidate the interaction of tau fibril (straight filament structure) at various lipid compositions using both all-atom and coarse grained MD simulations. The schematics of the straight filament structure of tau fibril along with the model bilayer are shown in Figure 1.

**Figure 1:**
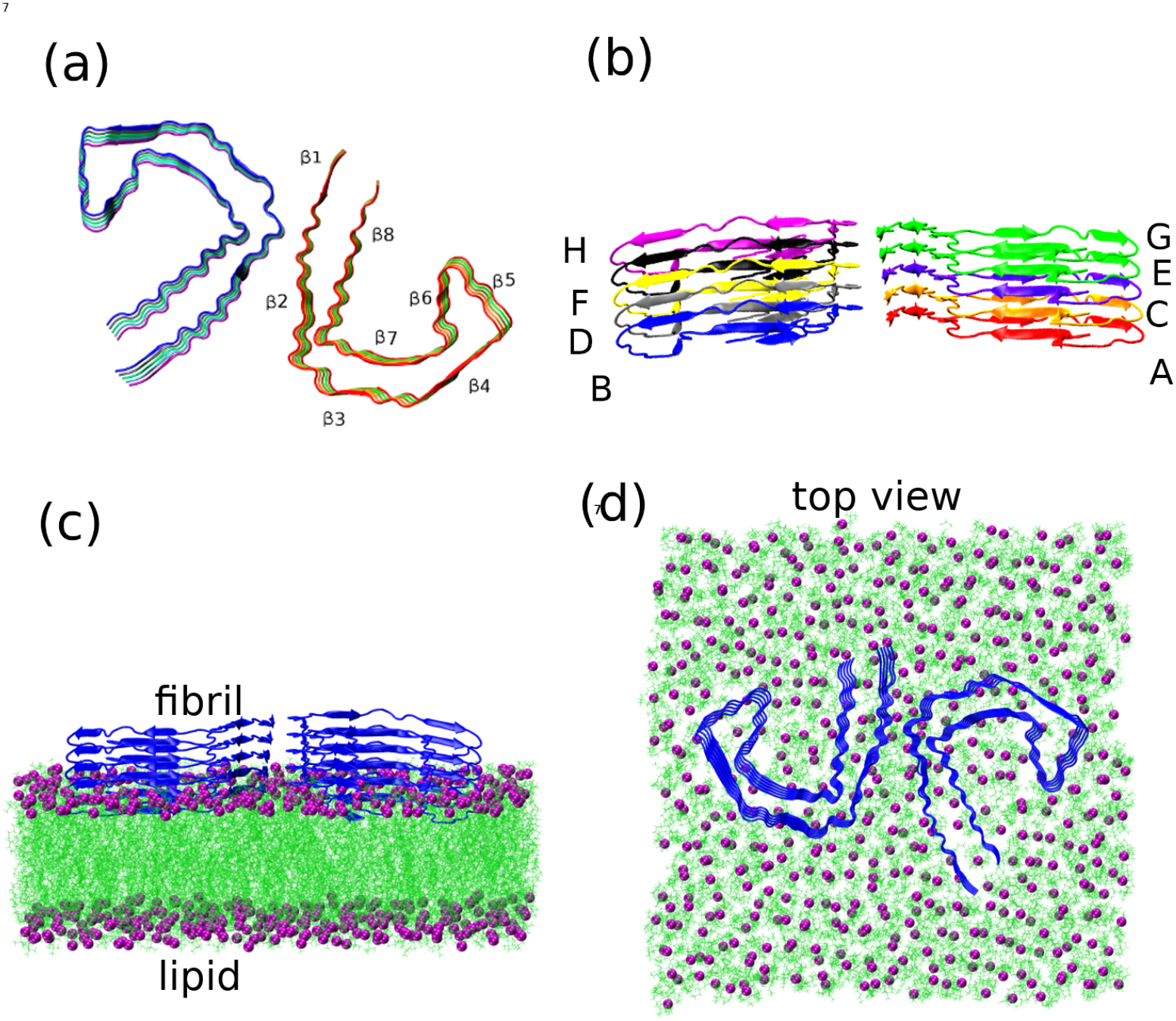
(a) The tau straight filament (SF) structure (PDB id- 503T) as seen from the top with each of the chains shown in different color. The strands of *β*-sheet regions are named sequentially from *β*1 to *β*8. (b) The individual chains of the tau-fibril are shown in different colors. (c) The fibril-lipid arrangement used in our simulations. The phosphorus atoms in the lipid head group are shown in purple, the lipid chains are shown in green and the fibril is shown in blue. (d) The fibril-lipid arrangement as viewed from the top.

## 2 Results and discussion

### 2.1 Area per lipid and Bilayer Thickness

We have modeled different systems to study the effect of the net charge of the lipid molecules and the effect of cholesterol on the tau fibril. Neutral POPC/POPE and negatively charged POPG are used to form the bilayer. In addition to that, a mixed composition of POPC and POPE is also used to build the bilayer. In this way seven compositions of the bilayer are included in our study. The schematics of the lipid molecules used in our study showing the sn-1 and sn-2 chains are given in the Supporting Information (Figure S1). Further details of the simulation setup is included in System Description. In the figures, the abbreviations PC, PC+CL, PE, PE+CL, PC+PE, PG, PG+CL are used to refer to POPC, POPC+CHOL, POPE, POPE+CHOL, POPC+POPE, POPG and POPG+CHOL lipid composition respectively. Cholesterol is known to induce changes in the properties of the bilayer. Cholesterol is necessary for the formation of lipid rafts and microdomains, and it adds stability to membrane proteins by increasing the membrane thickness and membrane lipid rigidity.^35^ The adsorption and the interaction of the tau proteins with the bilayers are the initial steps in the structural changes initiated over the cell membrane. Anionic lipids are proven to induce the tau aggregation.^10^ POPC consists of zwitterionic headgroups whereas the POPE consists of ethanolamine headgroup. POPG is an anionic lipid with a net charge of (−1). Negatively charged POPG lipids are found to influence the structure and conformation of other membrane proteins.^36^ To analyse the effect of tau peptide on the lipid membrane, we have computed the area per lipids (APL) and the membrane thickness at all the lipid compositions with and without the presence of the tau protein. The mean bilayer thickness and the area per lipids (APL) are given in Table 1. The area per lipid is calculated using the x and y dimensions of the box length and dividing the area by the total number of lipids in a leaflet. The bilayer thickness is computed using the average intra-phosphate distance in the bilayer. Figure 2 shows the change in bilayer properties depicted in a box and whisker plot. The presence of tau protein decreases both the APL and the bilayer thickness across all the lipid compositions. The significant decrease in the APL (0.06 nm^2^) is observed in the POPG, POPC+POPE and POPE+CHOL systems and the largest change (0.13 nm) in the membrane thickness is seen in the POPE systems.

**Table 1:** Variation of the bilayer properties with and without the tau protein.

| Bilayer | Area per Lipid<br>(nm<sup>2</sup>) | Bilayer thickness<br>(nm) |
| --- | --- | --- |
| POPC | 0.64 | 3.92 |
| POPC + tau | 0.60 | 3.88 |
| POPE | 0.56 | 4.28 |
| POPE + tau | 0.52 | 4.15 |
| POPG | 0.69 | 3.70 |
| POPG + tau | 0.63 | 3.64 |
| POPC + CHOL | 0.48 | 4.48 |
| POPC + CHOL + tau | 0.43 | 4.40 |
| POPE + CHOL | 0.46 | 4.56 |
| POPE + CHOL + tau | 0.40 | 4.53 |
| POPC + POPE | 0.60 | 4.11 |
| POPC + POPE + tau | 0.54 | 4.09 |
| POPG + CHOL | 0.50 | 4.29 |
| POPG + CHOL + tau | 0.45 | 4.23 |

**Figure 2:**
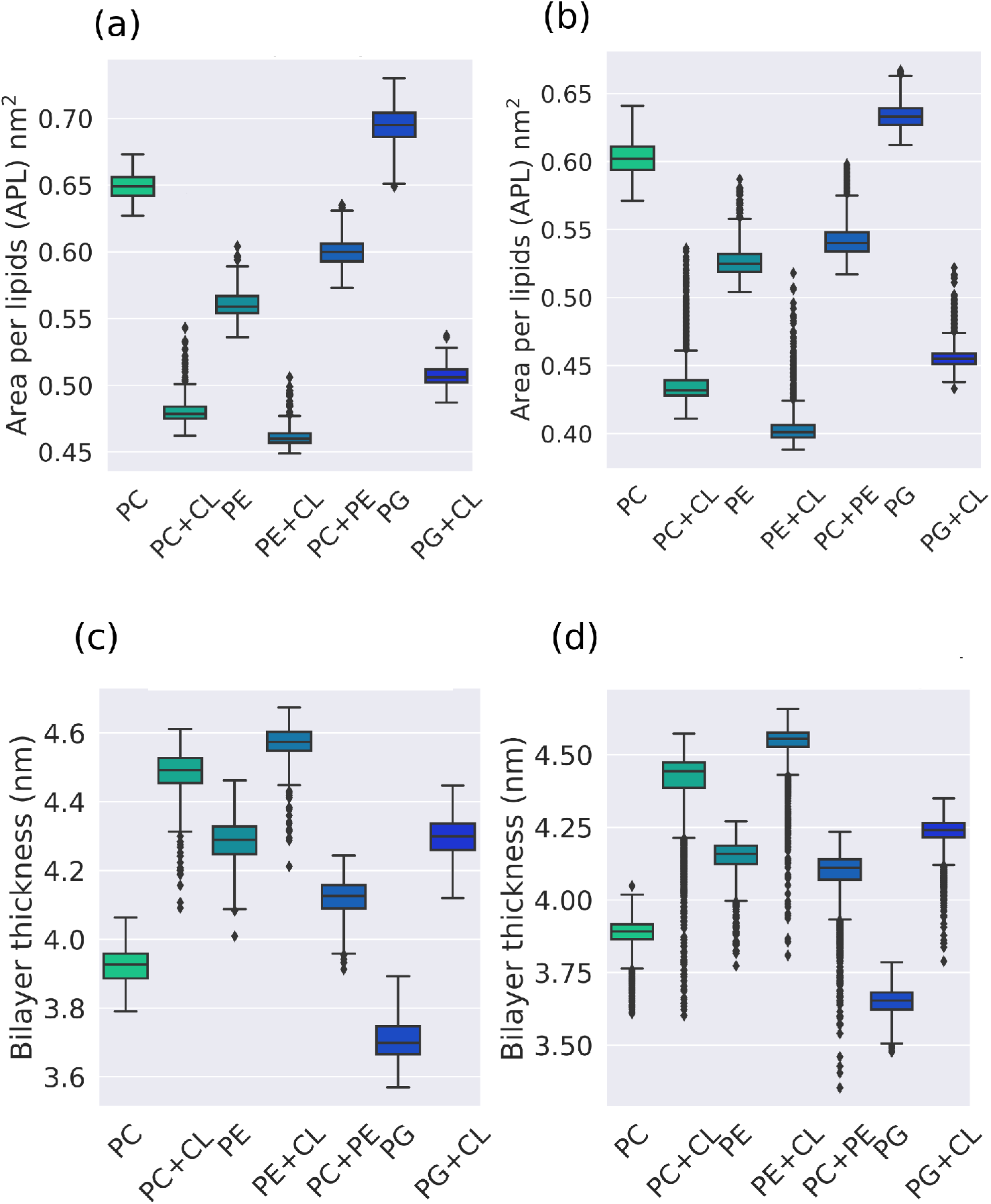
Box and whisker plots of the area per lipid and the bilayer thickness in case of the pure bilayers and bilayers with tau fibril. (a) and (b) Area per lipids (APL) for the pure lipids and the lipids with tau fibril respectively. (c) and(d) Bilayer thickness for the pure lipids and the lipids with tau fibril respectively.

To describe the local deformations of the lipid bilayers with the tau-protein, we have calculated the variation of the bilayer thickness in the system. The variation of bilayer thickness describing the local perturbations of the bilayer around the tau peptide in systems with POPG and POPG+CHOL in the presence of tau peptide is shown in Figure 3. The bilayer thickness for the other systems are given in the Supporting Information (Figure S3). The presence of CHOL is shown to increase the bilayer thickness in the region surrounding the tau-protein. As shown in the two cases of POPG and POPG+CHOL, it is found that the tau-protein deforms the bilayer surrounding the tau fibril. Hence, tau deform the negatively charged POPG membrane more than that of the POPG+CHOL membrane. The increased interaction between the negatively charged POPG bilayers and the tau-protein is also discussed in a subsequent section involving the number of contacts. A representative snapshot of a configuration of the POPG bilayer with the tau-protein is shown in Figure 3(c). The tau protein is seen to perforate the bilayer through its N-terminal region.

**Figure 3:**
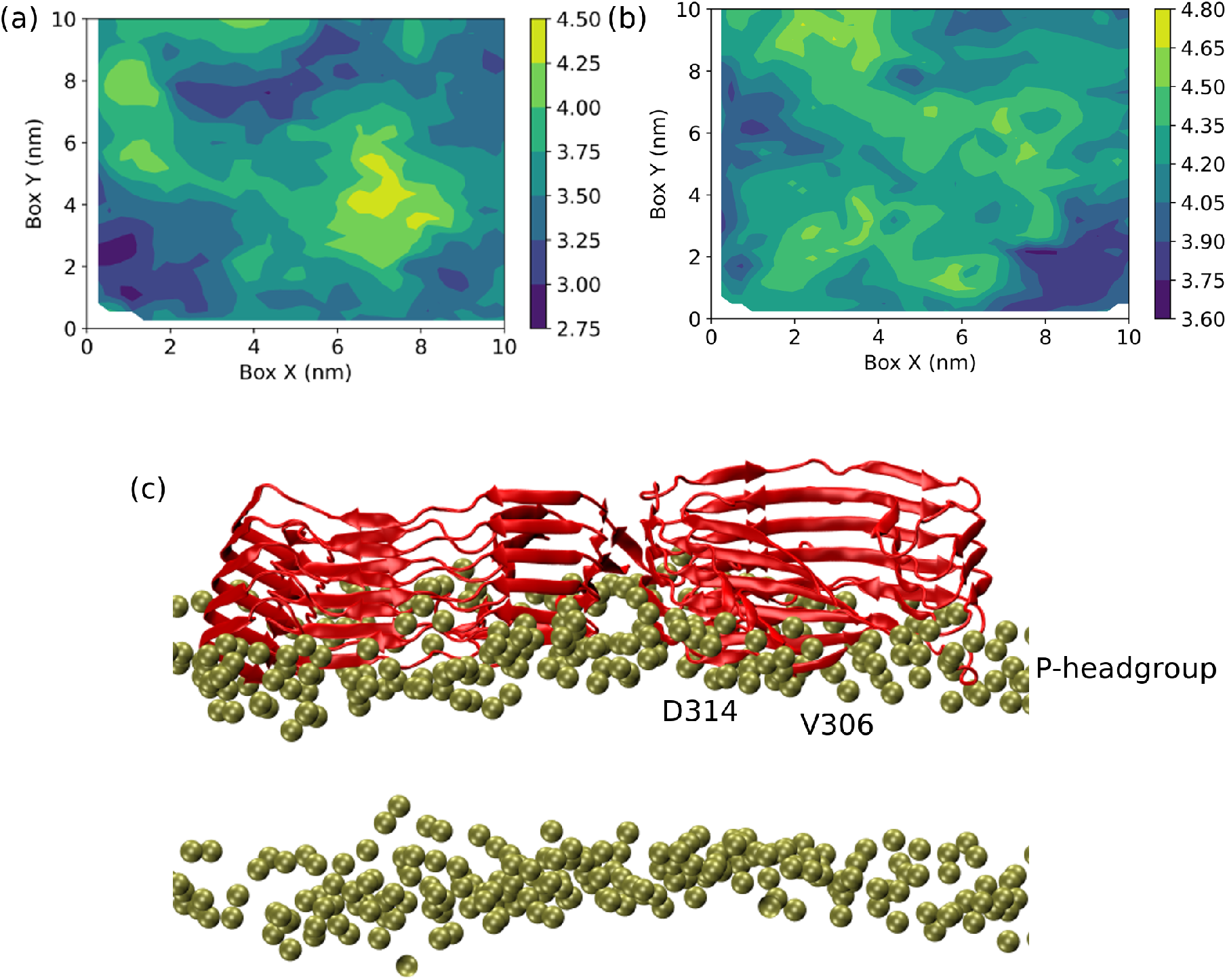
(a) and (b) are two dimensional thickness plots in the POPG and POPG+CHOL respectively, in the presence of tau-proteins. The thickness is shown in [nm] units in the color bar. (c) The representative snapshot of the tau-protein perforating the lipid bilayer in case of the POPG system. Only the head groups are shown for clarity.

### 2.2 Deuterium Order Parameter (S_*CD*_)

The order parameter (S_*CD*_) is important to quantify the structural deformation and flexibility of lipids in bilayers. Experimentally, the order parameters are derived from the NMR. The carbon atoms near the headgroup have higher order parameter, which decreases down the length of carbon chain since the movement of head groups are restricted and the tail regions are relatively free to move in the bilayer. S_*CD*_ is determined using the equation

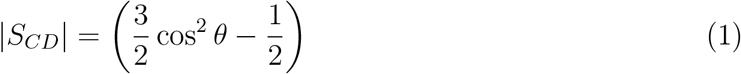

where *θ* is the angle between the C-H bonds in the lipid tail and the bilayer normal.

The presence of tau fibril leads to the decrease in the lipid order parameters as seen in case of POPE systems shown in Figure 4. The order parameters of the pure POPE lipids match with the earlier simulation and experimental results.^37–39^ The decrease in the order parameter is observed in both sn-1 and sn-2 chains. The decreasing S_*CD*_ order parameter correlates with the membrane thickness of the lipids in presence of tau protein and the pure lipids. The decreasing order parameter translates to the decreases in the bilayer thickness for lipid bilayers in presence of tau proteins. Higher order parameter indicate linear lipid chains which in turn leads to increase in bilayer thickness of the pure lipids.

**Figure 4:**
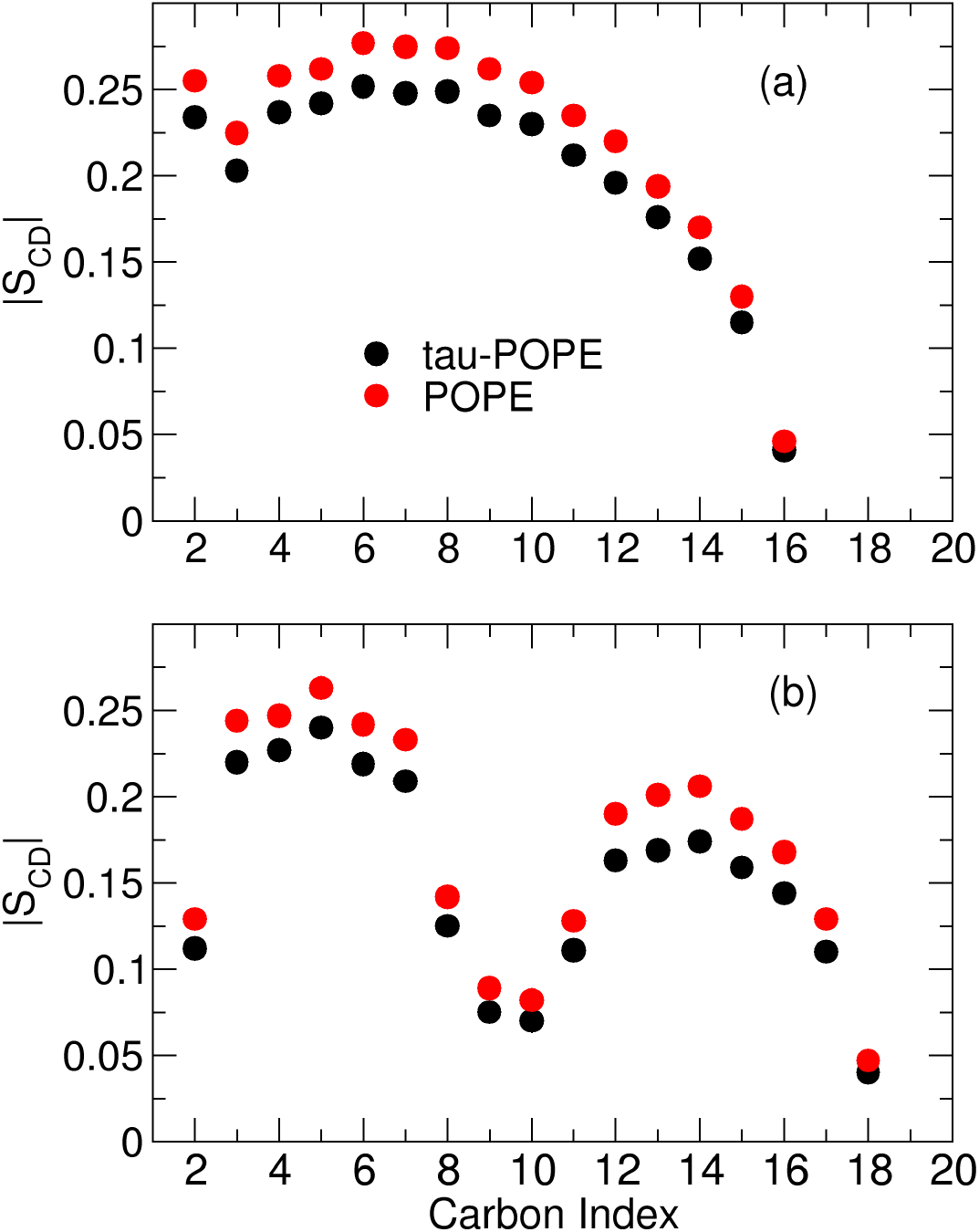
Deuterium order parameter (S_*CD*_) for the (a) sn-1 chains and (b) sn-2 chains in case of pure POPE and POPE with tau protein.

The addition of cholesterol in the bilayer leads to an increase in the membrane order parameters in case of both the pure lipids and lipids with tau, in accordance with the trend observed in the bilayer thickness. The order parameters for all the other systems are given in the Supporting Information (Figure S4 – S9).

### 2.3 Tau Fibril Stability

The structural differences of the tau fibril in various lipid membranes are studied using the root mean square deviations (RMSDs) and the root mean square fluctuations (RMSFs) of the tau protein. The RMSD and RMSF are calculated from the 400 ns of all-atom trajectory of the peptide – bilayer systems. The RMSD and RMSF plots are given in Figure 5. RMSDs show that the MD simulations are converged after 30 ns. The tau proteins in the POPE+CHOL and POPG+CHOL systems are the most stable as seen from the RMSF plots. Tau-peptides in the POPC+CHOL show the most fluctuations, followed by the those in pure POPC. Hence it is evident that the charge of the lipids and the incorporation of the cholesterol regulate the structural fluctuations of the tau peptide. The RMSD of tau peptide in water is observed to be around 0.5 nm (Figure S2, Supporting Information).

**Figure 5:**
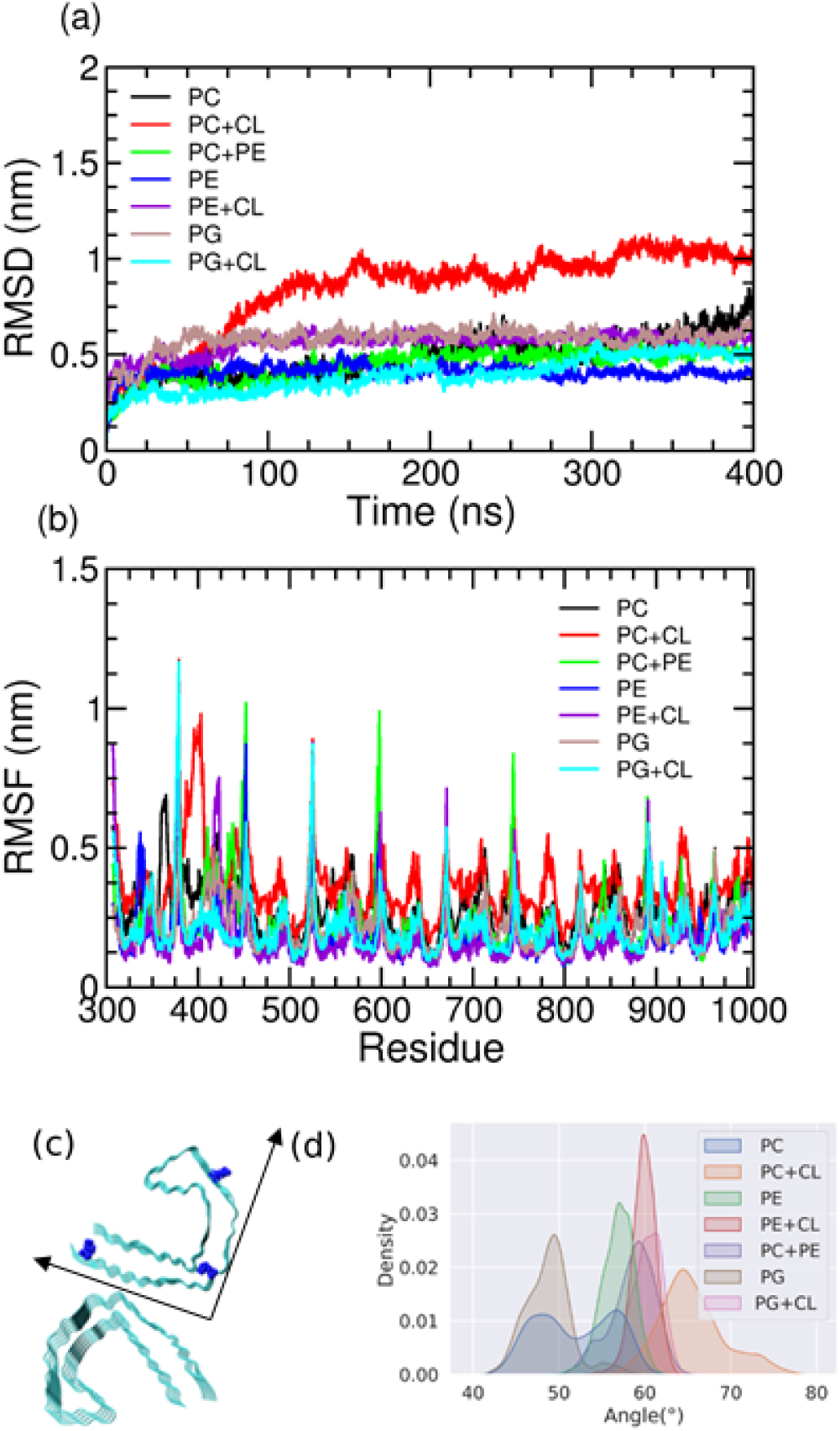
(a) RMSD of the C-*α* atoms of the tau peptides in the systems used in our study. (b) RMSF of the residues in the tau-peptide across all the systems (c) Pictorial representation of the angle of coverage used in the discussion. Three reference C-*α* residues are shown as blue VDW spheres. (d) The distribution of the angles in various bilayer systems.

To further analyze the dynamics of the tau proteins, we calculated the angle of coverage which is defined by the position of C-*α* residues as shown in the Figure 5(c). The angle of coverage defines the characteristic “C”-letter of the tau peptide from the cryo-EM structure. This angle is an important criterion to study the dynamics of the tau protofilament core and describes the area spanned by the tau-protein over the membrane plane. The distribution of this angle in all the studied systems are shown in Figure 5(d). We found that the angle of coverage depends on the lipid composition, and the incorporation of cholesterol increases the angle of coverage.

### 2.4 *β*-sheet content

Composition of the lipid membranes have an effect on the secondary structure content of the intrinsically disordered *β*-fibrils. The secondary structures of the intrinsically disordered proteins in turn effect their percolation within the lipid membranes. We have analysed the effect of the composition of the lipid membrane on the tau-structure. Previous MD simulations have given important insights into the secondary structures of amyloid fibrils interacting with lipid bilayer. ^40^ The distribution of the number of residues present as *β*-sheets are shown as violin plots in Figure 6. The negatively charged POPG lipids are found to increase the *β*-sheet content. Cholesterol decreases the *β*-sheet content of the tau fibril in case of POPC and POPE but increases in the case of POPG lipids. The mixed POPC – POPE lipids also increase the *β*-sheet content with respect to the pure POPE and POPC lipids. The number of *β*-sheet residues in case of the bulk water, with a mean value of 430 *±* 12, is higher than in presence of any lipids. The timeline of the secondary structures as evaluated from DSSP are shown in the Supporting Information (Figure S10-S13).

**Figure 6:**
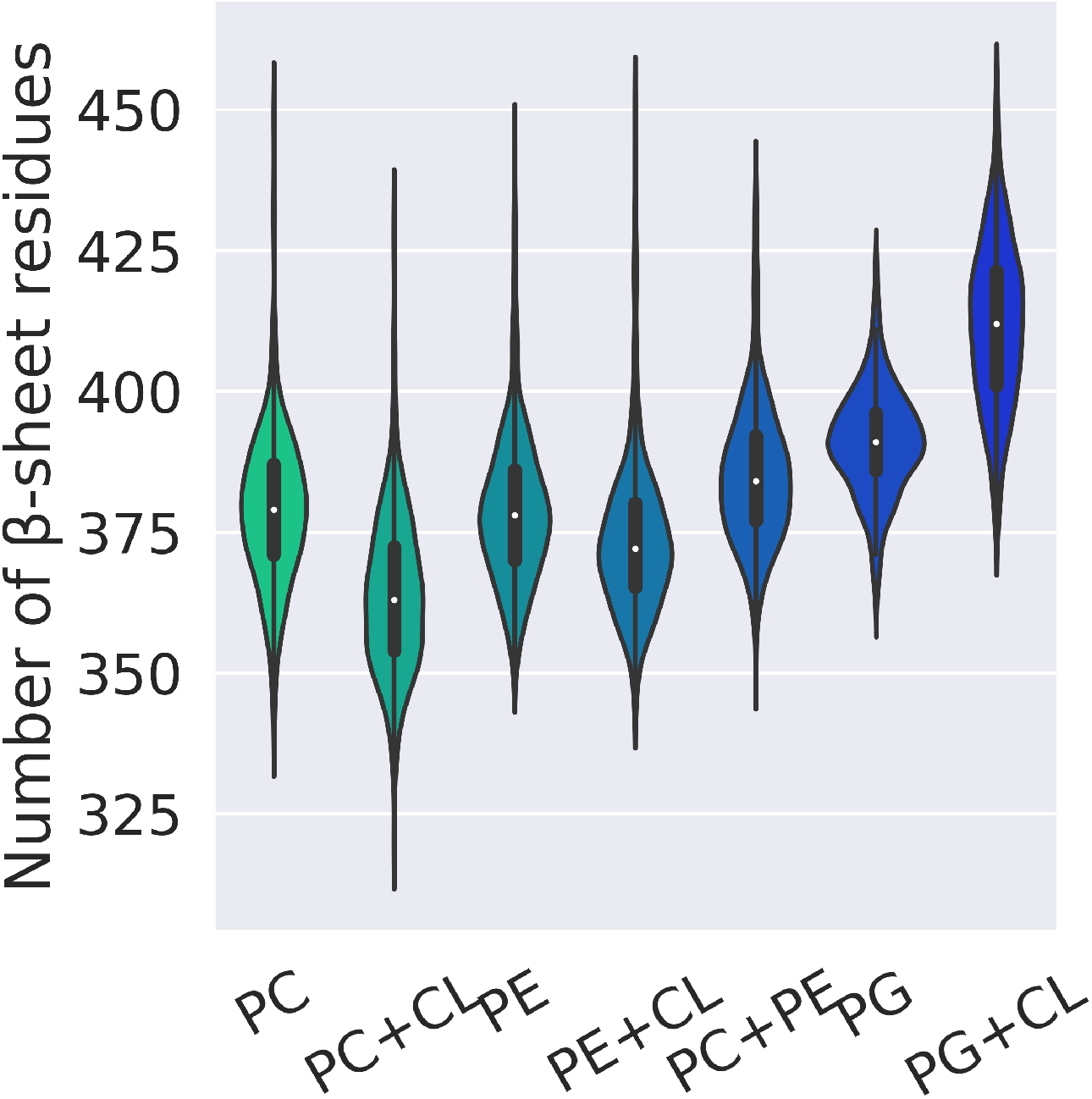
Violin plots of the distribution of the number of residues of tau protein present as *β*-sheet across all the lipid compositions.

The number of *β*-sheet residues over the simulation time is also shown in Supporting Information (Figure S14). It is evident that the presence of the charged membranes and the incorporation of the cholesterol modulate the *β*-sheet content in the tau fibrils.

### 2.5 Hydrogen Bonding

We have classified the hydrogen bonding according to the geometric criteria.^41^ In a strong hydrogen bond, the hydrogen atom and the acceptor are separated by a distance less than 2.2 Å, and the angle made by the donor, hydrogen atom, and the acceptor is within the range 130° – 180°. The corresponding distance and angular range are 2.0 – 3.0 Å and 90° – 180°, respectively, for a weak hydrogen bond. We found no strong hydrogen bonding in any of the trajectories. Inter- and intra molecular hydrogens bonds calculated from the simulations are useful in determining the structural properties of the bilayer at molecular level. The hydrogen bonding information of different systems (both intra- and inter-fibril) are given in Figure 7.

**Figure 7:**
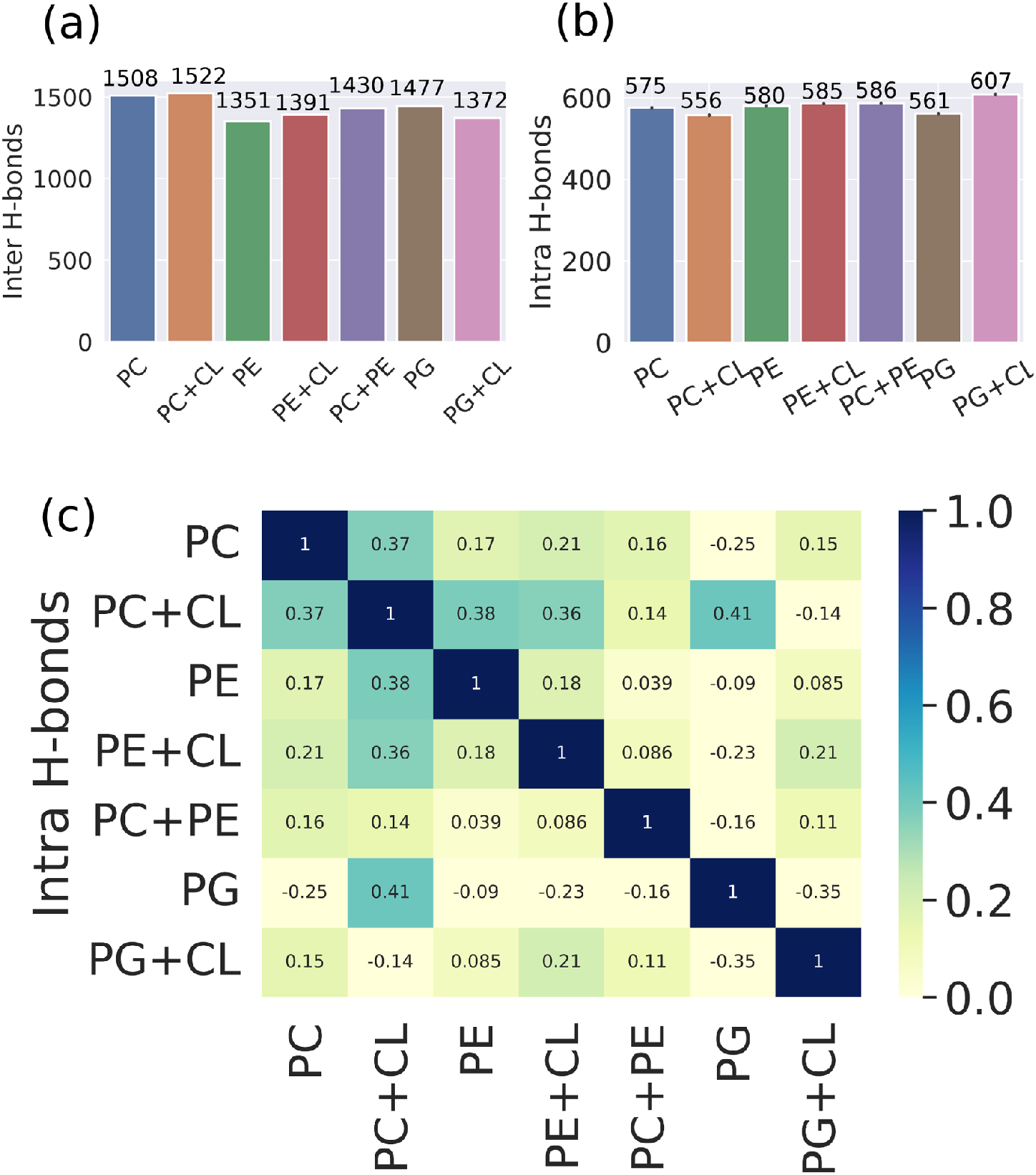
(a) The fibril-water hydrogen bonding and (b) intra-fibril hydrogen bonding across all the systems. (b) The probability to form intra-fibril hydrogen bonding shown through a correlation matrix for all the lipids. The scale for the magnitude of the correlation is shown in a color bar.

The probability of intra-fibril hydrogen bonding is found to correlate with the charge of the lipid and the presence of cholesterol as shown in the correlation matrix in Figure 7(c). Correlation matrix is an easy way to summarize the correlation between all the variables in a dataset. It is calculated through the Pearson correlation coefficient with -1 meaning a negative correlation, +1 meaning a positive correlation and 0 meaning no correlation. Each element of the matrix is the correlation coefficient (c_*ij*_) of the intra-fibril hydrogen bonding across the systems. The correlation matrix is defined as follows

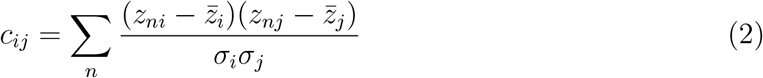

where c_*ij*_ denotes the *ij*-th element of the correlation matrix, (*i, j*) are the two different sets of data points across which the correlation is calculated, *z*_*ni*_ is the number of intra-fibril hydrogen bonds in the n-th frame of the system *i*, 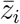 and 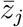 are the mean number of intra-fibril hydrogen bonds over all the frames in the system and *σ*_*i*_, *σ*_*j*_ are the standard deviations of intra-fibril hydrogen bonds in the system *i, j* respectively.

The negatively charged POPG and POPG+CHOL are inversely correlated whereas the systems with POPC/POPE lipids are directly correlated with POPC+CHOL/POPE+CHOL. The trend is similar to the variation of the number of residues of tau present as *β*-sheets in the bilayer with cholesterol, as the intra-fibril hydrogen bonds keep the *β*-sheet structure of the tau protein. In presence of cholesterol, *β*-sheets are stabilized in negatively charged POPG+CHOL system due to the increase in the number of intra-fibril hydrogen bonds.

For the intermolecular hydrogen bonding between the tau and water, no clear trend is observed. The cholesterol increases the tau-water hydrogen bonding in the POPE, decreases the hydrogen bonding in POPG and shows negligible difference in the POPC.

### 2.6 Number of Contacts

We calculated the number of lipid contacts with peptide to characterize the peptide–lipid interaction. The timeline for the number of contacts are given in Figure 8(a). The coarse grained (CG) trajectories are used to model the long timescales of the binding of proteins to the lipids. The CG cutoff for the contacts are taken to be 0.7 nm according to the *α*-synuclein binding work by Sansom group.^42^ The center-of-mass distance between the tau-peptides and the lipids are computed across all the trajectories and are given in the Supporting Information (Figure S15 – S16). The highest number of contacts are observed in the system with POPG lipids, and the least number of contacts in system with POPC lipids. Cholesterol increases the number of contacts in both zwitterionic and anionic lipids. The number of contacts is reduced in mixed POPC+POPE and is comparable to the POPC lipids. Thus the negatively charged lipids increase the binding affinity of the tau-peptides to the membrane. The principal structures obtained by the clustering algorithm (shown in Figure 8) also reveal that the tau-peptides show differential mode of interaction with the lipids.

**Figure 8:**
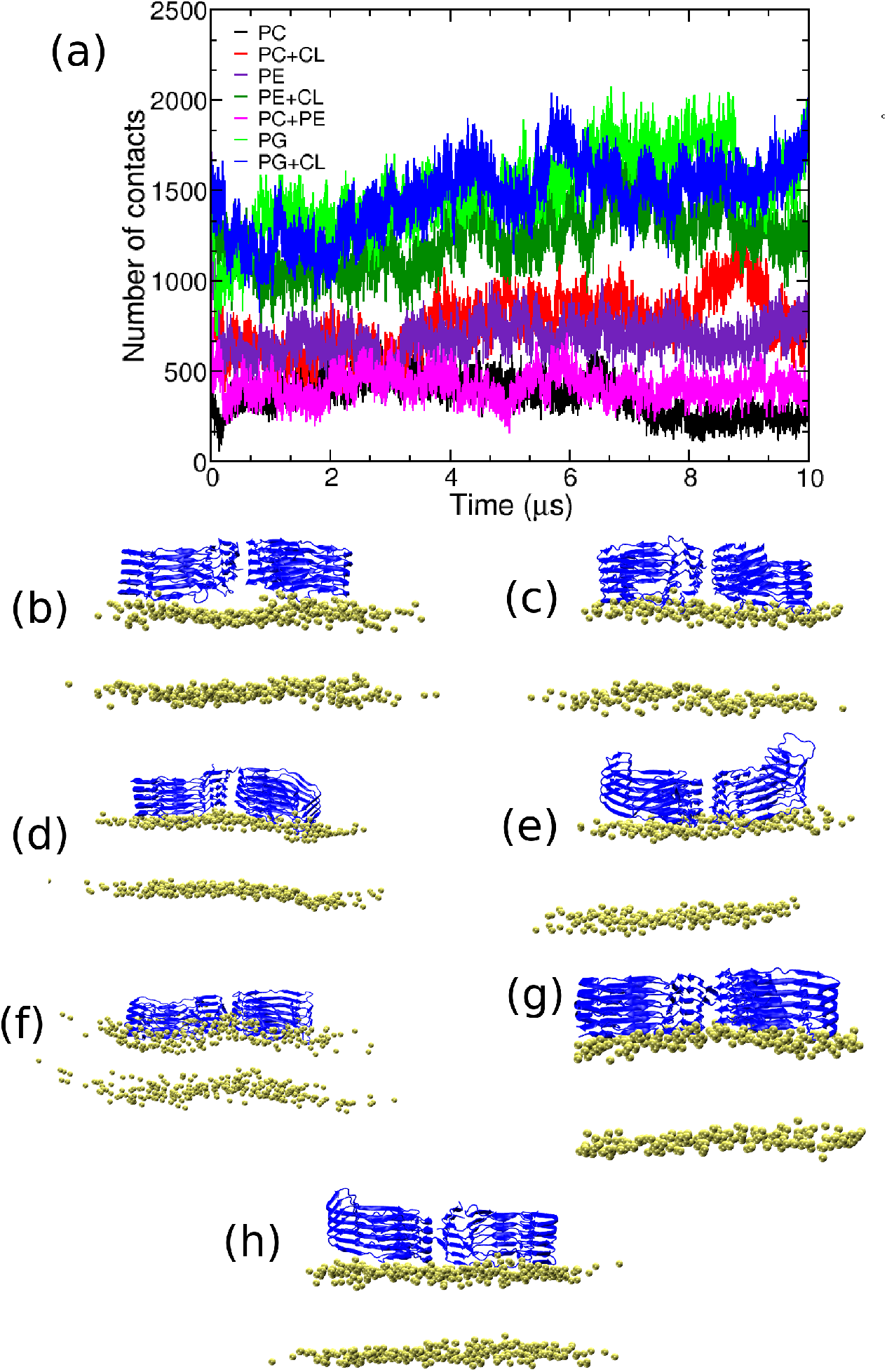
(a) Number of peptide – lipid contacts across all the trajectories. (b)-(h) are the most probable structures obtained through the clustering algorithm. The peptide is shown in new cartoon representation in color blue and the phosphorus atoms are shown as VDW spheres.

## 3 Conclusion

The knowledge of the interaction of the tau proteins with the lipid membranes are pivotal to the study of the effect of tau peptides on the cell membrane. The detailed molecular level interaction of the tau proteins with lipids and their effect in the tauopathy is still unavailable. This is due to the complex lipid compositions and the difficulty in characterizing the tau-peptides experimentally. The MD simulations of the tau-peptides in seven different lipid systems, comprising of zwitterionic and charged lipids have been carried out. The tau proteins are found to preferentially bind to anionic lipids than to the neutral lipids in agreement with the experimental reports.^9^ The addition of cholesterol leads to an increase in the thickness of the bilayer and the lipid tail order, but decreases the area per lipid. Cholesterol infused lipids in our model bilayers induce definite changes in the tau-conformational states. The changes induced by the cholesterol also differs between the anionic POPG and the zwitterionic POPC/POPE membranes. Hence, our study illustrate that the composition of lipids in the cell-membrane influences the interaction and binding of tau-peptides.

## 4 Methodology and simulation details

### 4.1 Coarse-grained simulations

The protein structure of the pdb id – 503T was coarse grained using the Martini representation. Martini version 2.0 and 2.2 have been used to model the lipids and proteins respectively.^43,44^ CHARMM-GUI was used to generate the initial configurations for the fibril and lipids.^45^ The simulations were performed using the parallelized GROMACS molecular dynamics code (version 5.1.4).^46^ The system was equilibrated after the initial energy minimization. The initial equilibration was done for the systems in six steps, restraining the proteins and lipids sequentially, according to the CHARMM-GUI protocol. Electrostatic and Lennard-Jones interactions were modeled up to 1.1 nm. The system was coupled to a thermostat at 310 K with a coupling constant of 1.0 ps using the v-rescale thermostat. ^47^ Pressure was maintained at 1 bar with a coupling constant of 5 ps using the semi-isotropic Berensden algorithm.^48^ Production runs were performed using a time step of 20 fs for a duration of 10 *μ*s. Electrostatic interactions were modeled using the reaction field method using dielectric constant of 15, following the recommended simulation parameters for coarse-grained Martini. The potential shift Verlet scheme was used for the Lennard-Jones cutoff at long distances. The initial velocities for the systems were chosen from a Maxwell distribution at 310 K. Final production runs were performed using the v-rescale thermostat and Parrinello-Rahman barostat with coupling constants of 1 ps and 12 ps respectively.^49^ The details of the studied systems are presented in Table 2.

**Table 2:** Details of the fibril and lipid systems used in the study.

| System description | Number of lipids | Number of waters | Production run (ns) | Number of atoms |
| --- | --- | --- | --- | --- |
| All-atom (protein and lipids) |  |  |  |  |
| POPC | 510 | 46400 | 400 | 219248 |
| POPE | 510 | 40692 | 400 | 197500 |
| POPG | 510 | 42358 | 400 | 203938 |
| POPC + CHOL (7:3) | 510 | 41478 | 400 | 195424 |
| POPE + CHOL (7:3) | 510 | 31334 | 400 | 161569 |
| POPG + CHOL (7:3) | 550 | 42463 | 400 | 200461 |
| POPC + POPE (1:1) | 510 | 44309 | 400 | 210666 |
| protein (no-lipids) | — | 106819 | 40 | 332515 |
| All-atom (pure lipids) |  |  |  |  |
| System description | Number of lipids | Number of waters | Production run (ns) | Number of atoms |
| POPC | 200 | 12300 | 40 | 63764 |
| POPE | 200 | 10856 | 40 | 56812 |
| POPG | 200 | 11116 | 40 | 59004 |
| POPC + CHOL (7:3) | 200 | 10819 | 40 | 55713 |
| POPE + CHOL (7:3) | 200 | 9652 | 40 | 50946 |
| POPG + CHOL (7:3) | 200 | 10040 | 40 | 52532 |
| POPC + POPE (1:1) | 200 | 11488 | 40 | 60422 |
| Coarse-grained (protein and lipids) |  |  |  |  |
| System description | Number of lipids | Number of water beads | Production run (μs) | Number of beads |
| POPC | 510 | 12078 | 10 | 20162 |
| POPE | 510 | 9822 | 10 | 17846 |
| POPG | 510 | 10410 | 10 | 18864 |
| POPC + CHOL (7:3) | 510 | 10309 | 10 | 17739 |
| POPE + CHOL (7:3) | 510 | 9697 | 10 | 17468 |
| POPG + CHOL (7:3) | 550 | 10132 | 10 | 18273 |
| POPC + POPE (1:1) | 510 | 11154 | 10 | 19214 |

### 4.2 All-atom simulations

The initial structure of the fibril was taken from the pdb id – 503T for the all atom simulations with CHARMM-36m parameters. ^22^ The fibril was placed in the pre-equilibrated bilayer using the CHARMM-GUI module.^45^ Water and chloride ions were added to solvate and neutralize the system. TIP3P was used to model the water molecules.^50^ Additional 0.15(M) of KCl was added to maintain the physiological salt concentration. The lipid molecules and cholesterol were modeled using the CHARMM-36 parameters by Klauda et al. ^51,52^ Energy minimization was performed to avoid steric clashes. The temperature was maintained at 310K using the Nosé-Hoover thermostat^53^ with the coupling constant of 1.0 ps. The equilibration of the lipid bilayers were performed at subsequent steps using the CHARMM-GUI protocol by applying restraints on the proteins and lipids. The final production runs were performed in the NPT ensemble for 400 ns using a time step of 2 fs. The pressure was maintained at 1 bar using the semi-isotropic Parrinello-Rahman barostat.^49^ The long range electrostatics were treated using the particle mesh Ewald (PME).^54^ Lennard-Jones interactions were calculated up to a cutoff distance of 1.2 nm with a force-switch function. The bonds involving hydrogen atoms were constrained using the LINCS algorithm.^55^ All-atom simulations were carried out using the GROMACS (version 5.1.4) MD code.^46^

The analyses have been done using the in-house codes and GROMACS tools. The *β*-sheet content was analyzed using the DSSP utility in GROMACS. Clustering analysis was done using the Gromos algorithm implemented by Daura *et. al*^56^ with the RMSD cutoff of 0.2 nm.

### 4.3 System Description

We have simulated 14 systems comprising of 1-palmitoyl-2-oleoyl-sn-glycero-3-phosphocholine (POPC), 1-palmitoyl-2-oleoyl-sn-glycero-3-phosphatidylethanolamine (POPE), 1-palmitoyl-2-oleoyl-sn-glycero-3-phosphatidylglycerol (POPG) along with cholesterol (CHOL) at seven different compositions. Additionally, we have simulated the tau protein in a water box. Throughout our study, we have used symmetric composition of lipids in the upper and the lower leaflet of the bilayer. In case of the pure POPC system 255 POPC molecules are randomly placed both on the upper and lower leaflet with the tau peptide being placed on the membrane surface. In case of the POPC/POPE+CHOL systems, 179 POPC/POPE + 77 CHOL molecules are randomly placed on the upper leaflet and 178 POPC/POPE + 76 CHOL molecules are placed on the lower leaflet and are then packed by placing the tau-peptide over the membrane surface. In POPG+CHOL, the upper leaflet comprises of 193 POPG and 83 CHOL, and the lower leaflet comprises of 192 POPG and 82 CHOL. Finally, in the POPC+POPE systems the upper and the lower leaflet comprises of 128 POPC + 127 POPE molecules respectively along with the tau peptides. The simulation details are given in Table 2.

## Acknowledgement

The authors gratefully acknowledge NISER Bhubaneswar for the computational resources.

## Supporting Information Available

The supporting information comprises of the schematics of the lipid molecules used in our study (Figure S1), RMSD of tau-peptide in the water box (Figure S2), bilayer thickness projected over the bilayer plane (Figure S3), order parameters (S_*CD*_) of the lipid tails (Figure S4-S9), DSSP timelines for all the systems (Figure S10-S13), number of beta-sheet residues (Figure S14) and distance profiles of the CG-models (Figure S15-S16).

## Table of Contents Graphic

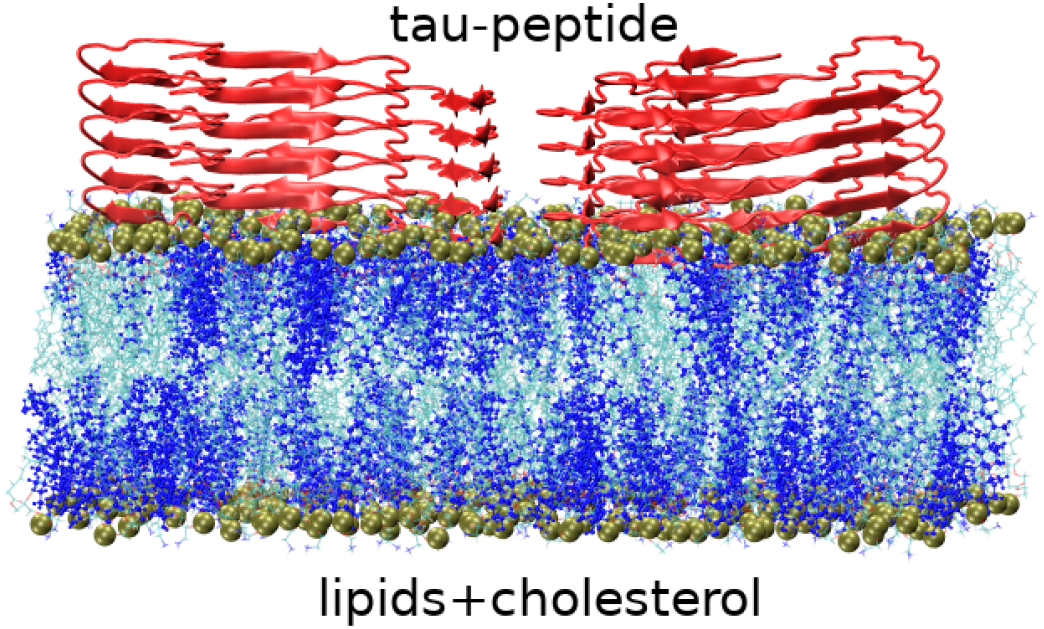

## Notes

### Competing Interest Statement

The authors have declared no competing interest.

