## Supporting information for "The Effect of Lipid Composition on the Dynamics of Tau Fibrils"

#### Contents

**1 Structure of lipids** **2**

**2 RMSD** **3**

**3 Bilayer thickness projected on the bilayer plane** **4**

**4 Acyl Chain Order Parameter ($S_{CD}$)** **5**

**5 DSSP timelines across all the systems** **11**

**6 Number of $\beta$-sheet residues** **15**

**7 Distance Profiles for the CG models** **16**

### 1 Structure of lipids

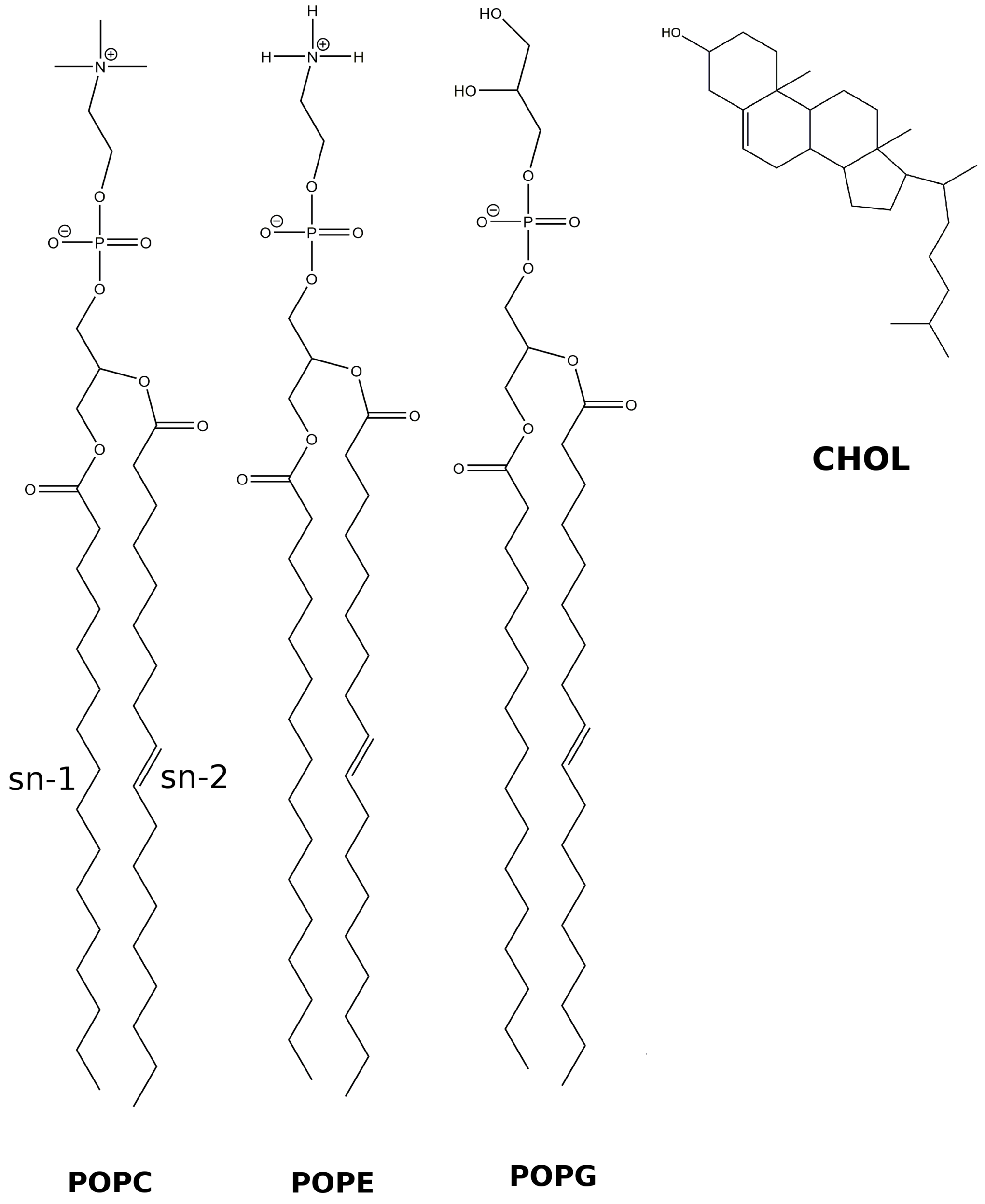


Figure S1: Schematics of the lipid molecules used in our study.

#### 2 RMSD

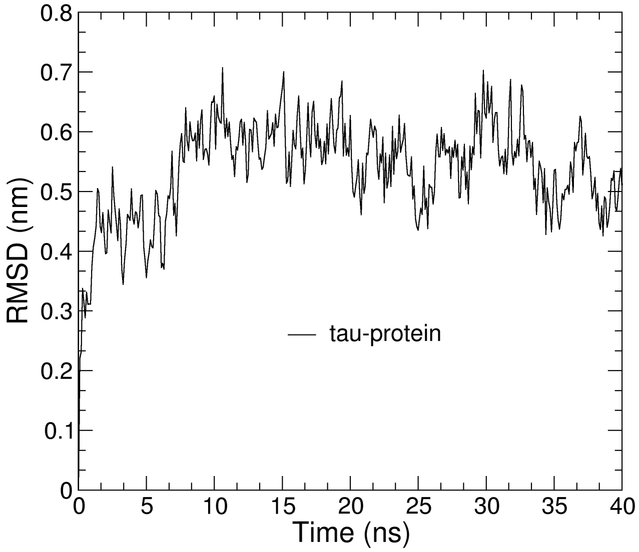


Figure S2: The RMSD for the tau-peptide in the water box.

### 3 Bilayer thickness projected on the bilayer plane

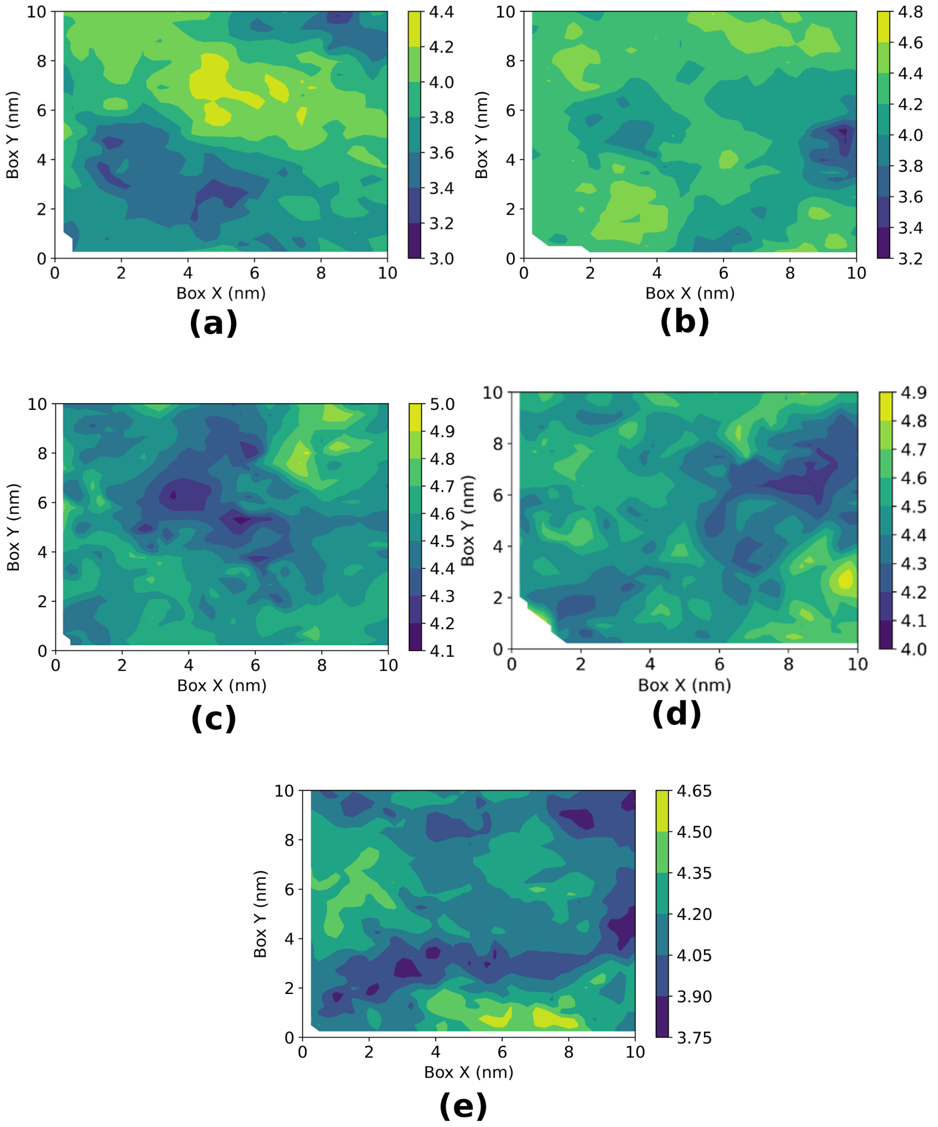


Figure S3: The bilayer thickness for (a) POPC (b) POPE (c) POPE+CHOL (d) POPC+CHOL (e) POPC+POPE systems calculated over the bilayer plane. The thickness is shown in [nm] units in a colorbar.

### 4 Acyl Chain Order Parameter ($S_{CD}$)

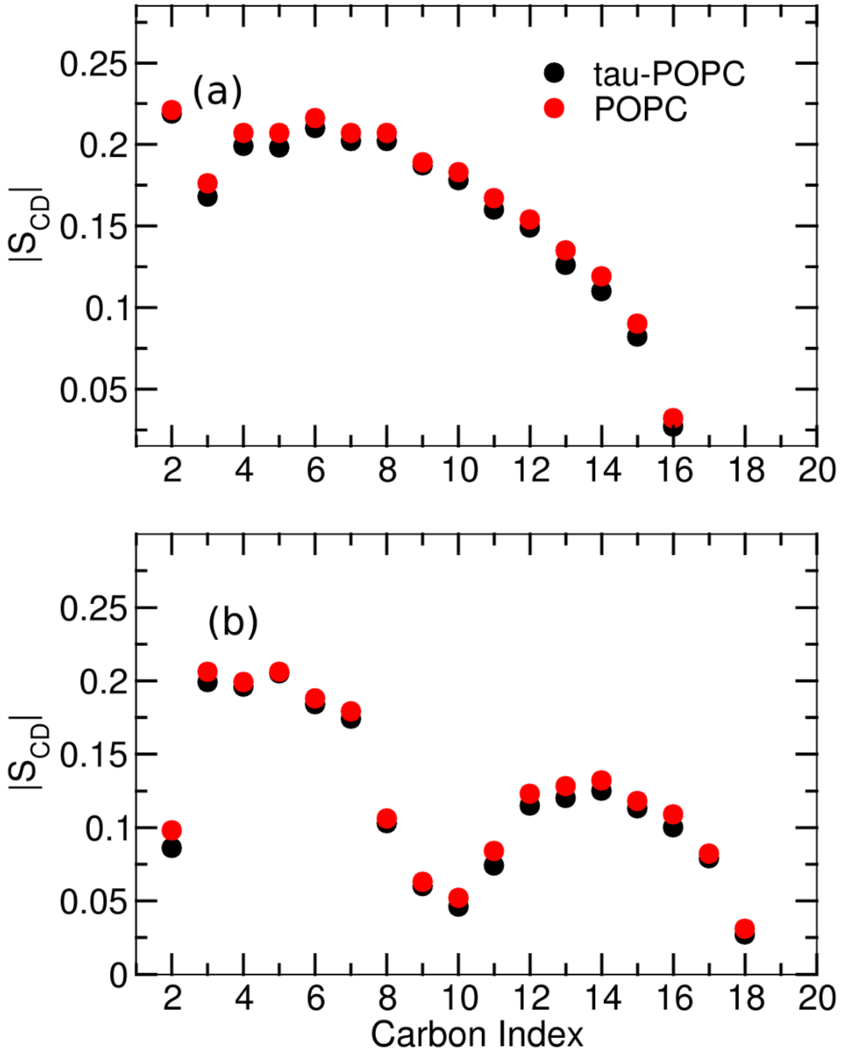


Figure S4: Deuterium order parameter ($S_{CD}$) for the (a) sn1 chains and (b) sn2 chains respectively.

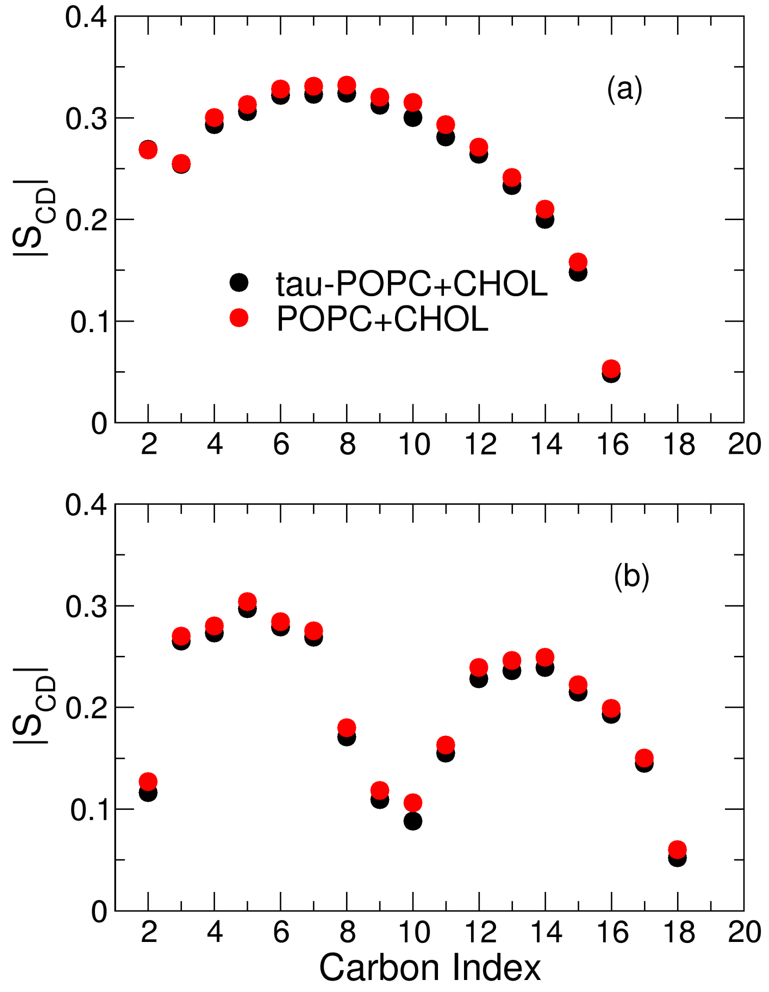


Figure S5: Deuterium order parameters for the (a) sn-1 chains (b) and for the sn-2 chains respectively.

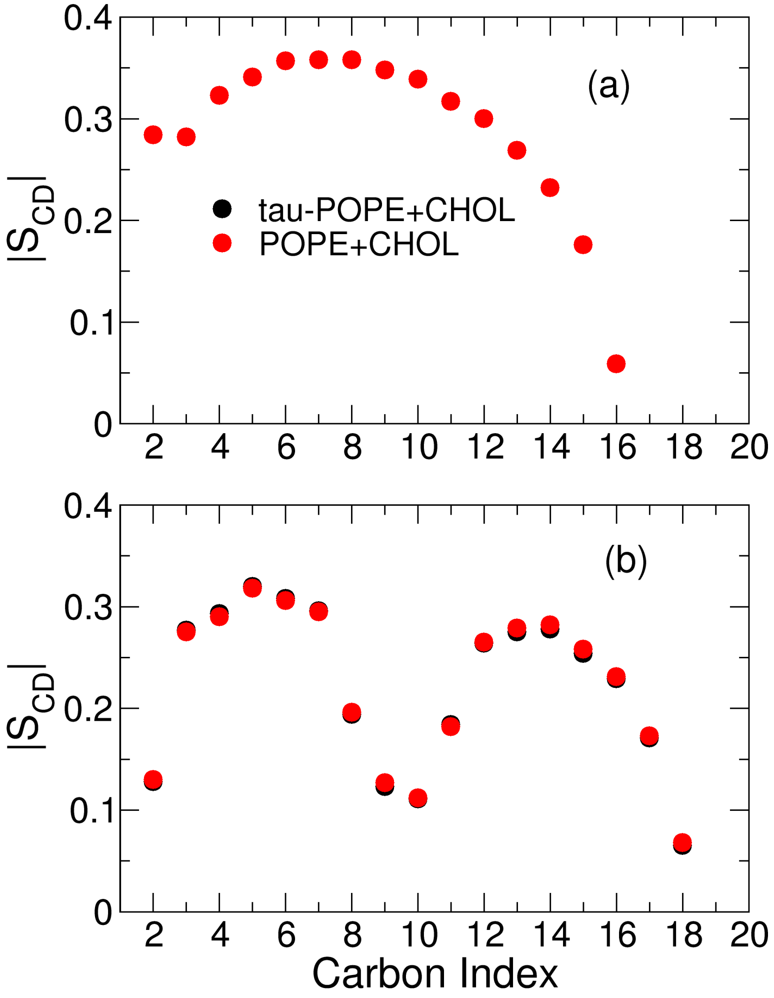


Figure S6: Deuterium order parameters for the (a) sn-1 chains (b) and for the sn-2 chains respectively.

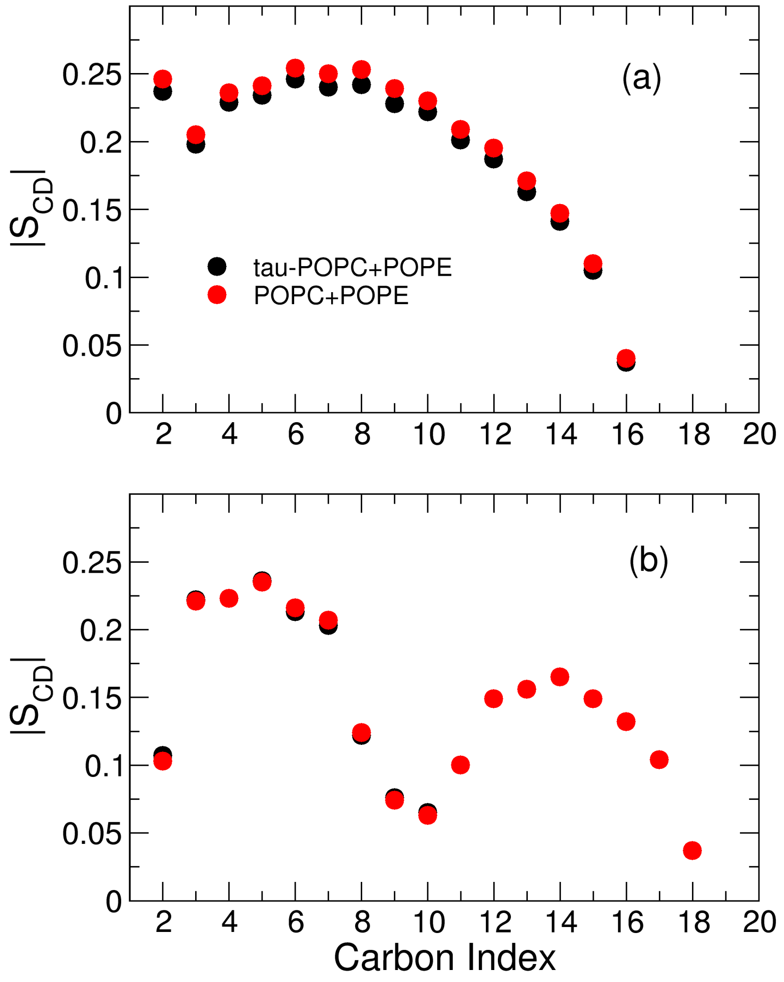


Figure S7: Deuterium order parameters for the (a) sn-1 chains (b) and for the sn-2 chains respectively.

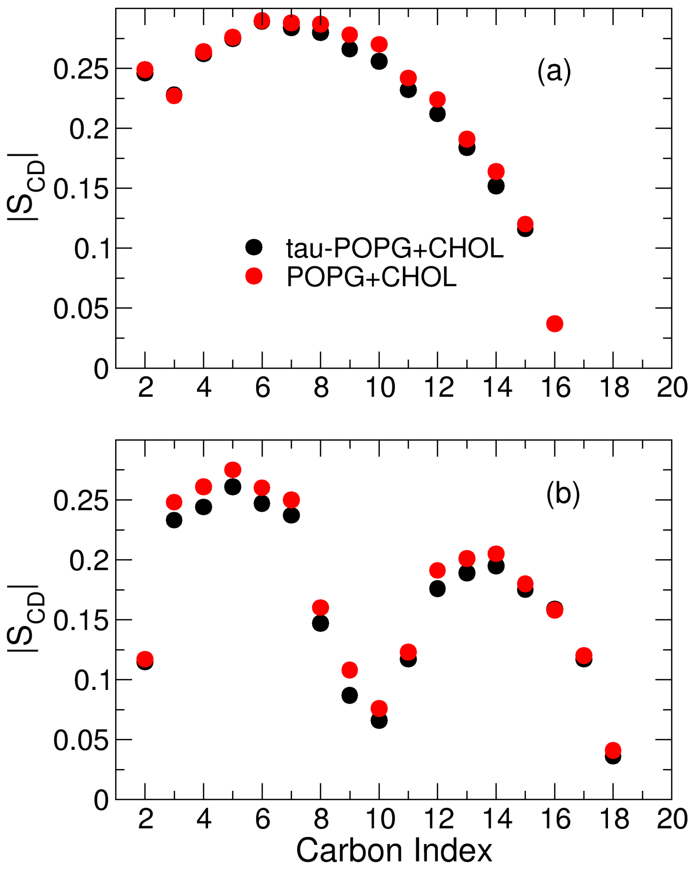


Figure S8: Deuterium order parameters for the (a) sn-1 chains (b) and for the sn-2 chains respectively.

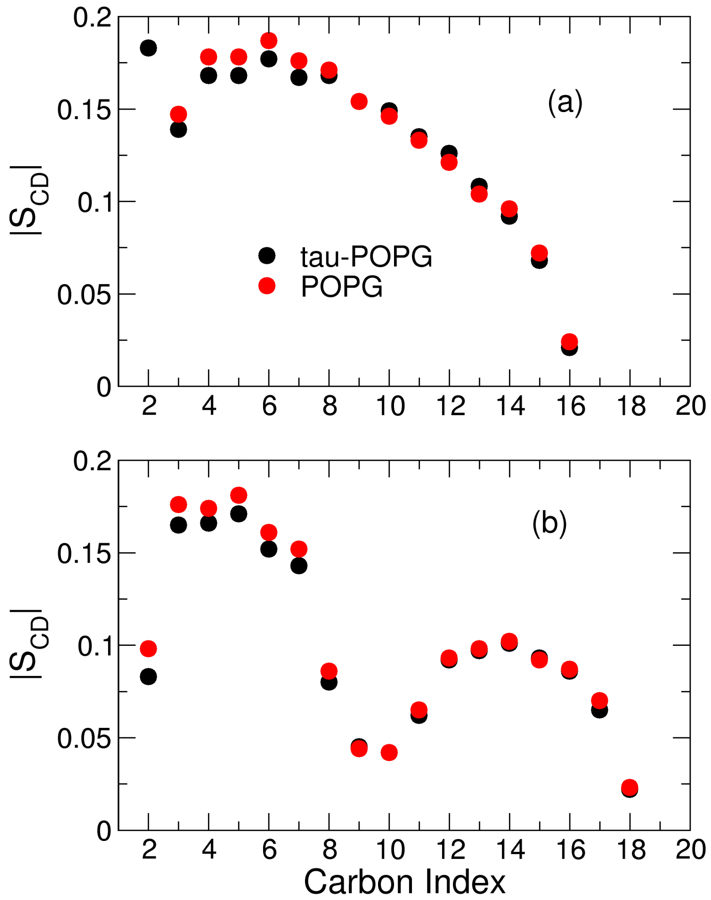


Figure S9: Deuterium order parameters for the (a) sn-1 chains (b) and for the sn-2 chains respectively.

### 5 DSSP timelines across all the systems

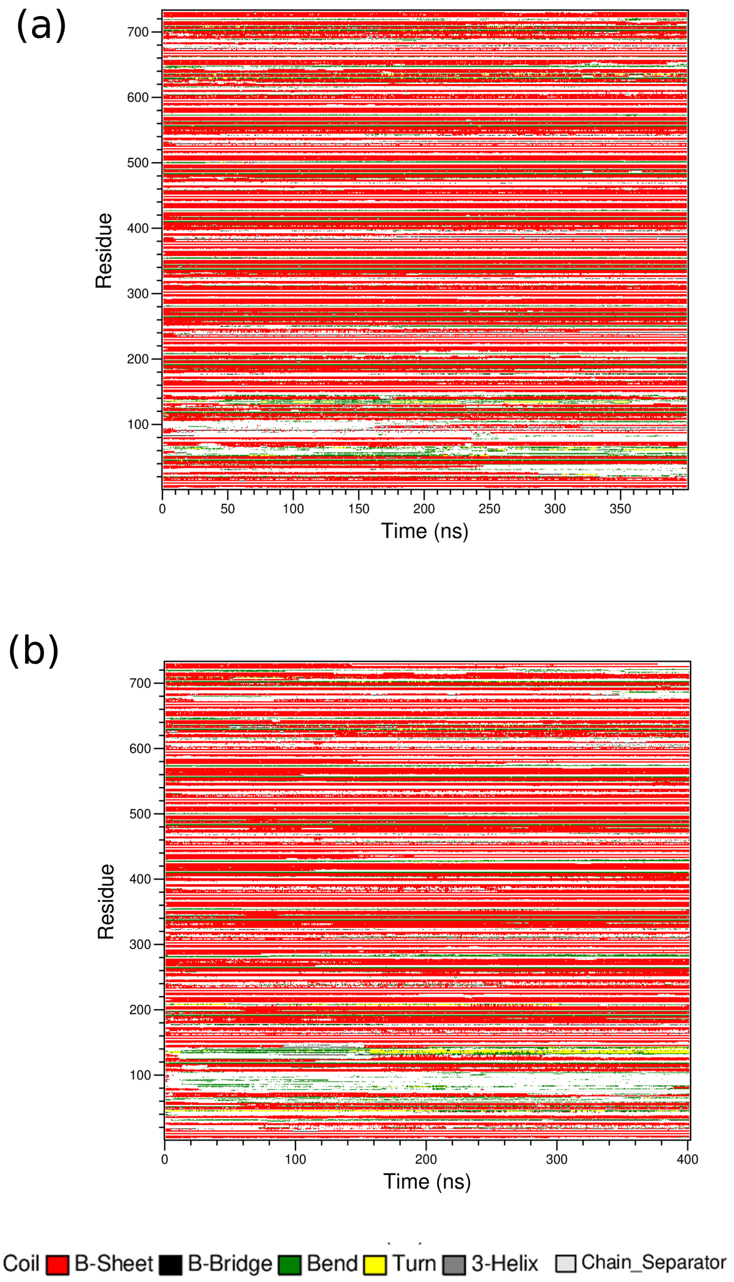


Figure S10: The DSSP for the systems (a) POPC (b) POPC+CHOL.

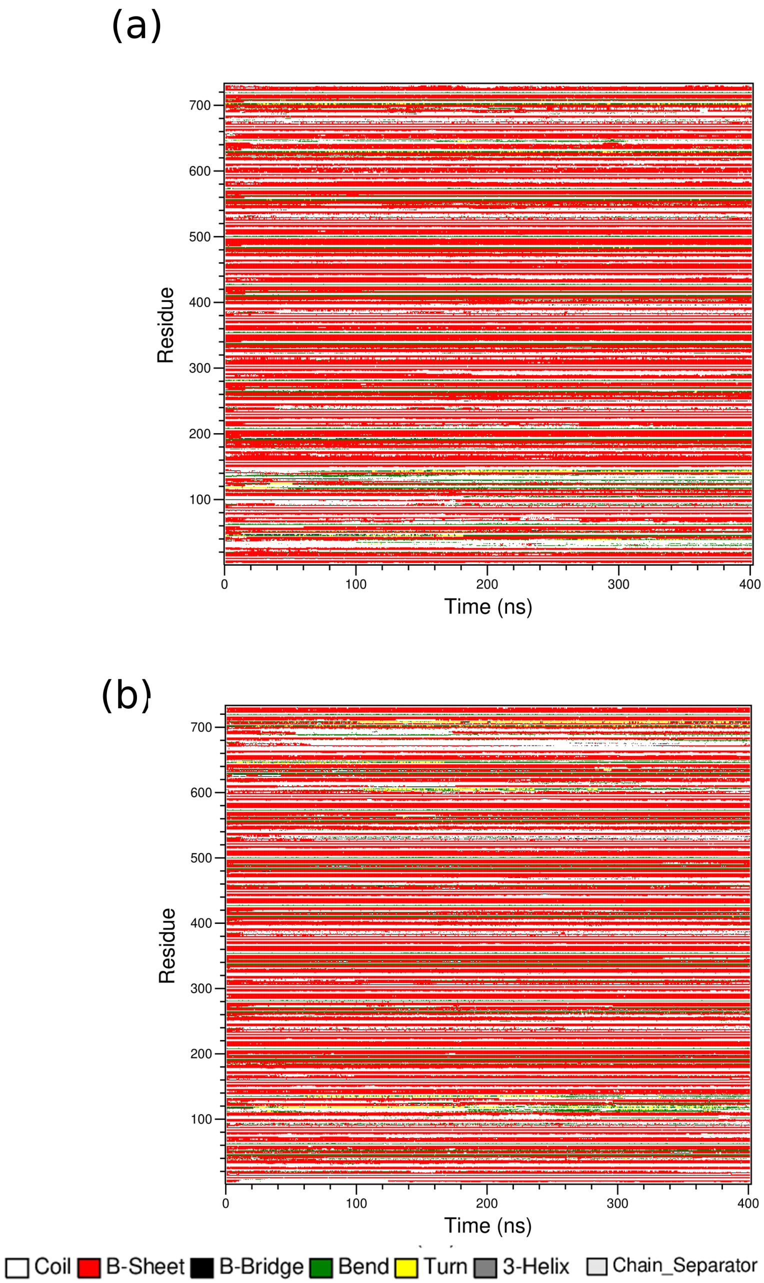


Figure S11: DSSP timelines for the systems (a) POPE (b) POPE+CHOL.

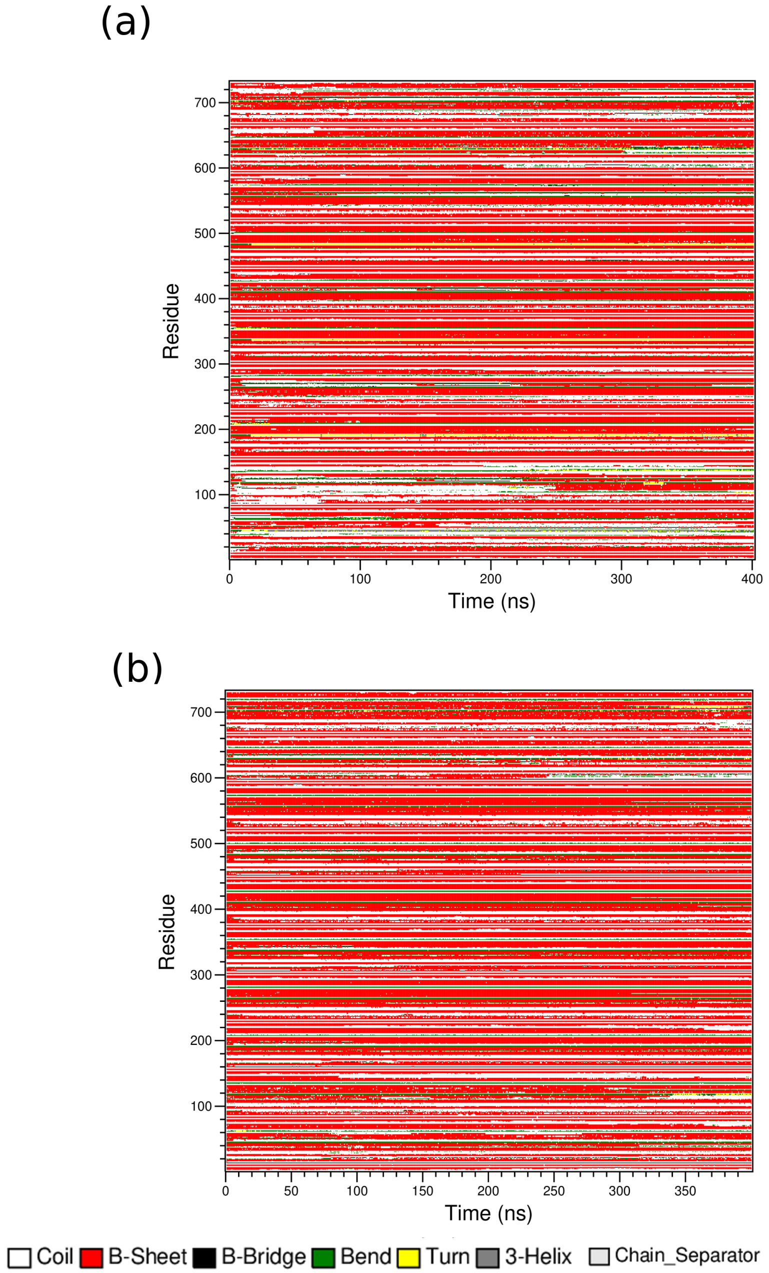


Figure S12: DSSP timelines for the systems (a) POPG (b) POPG+CHOL.

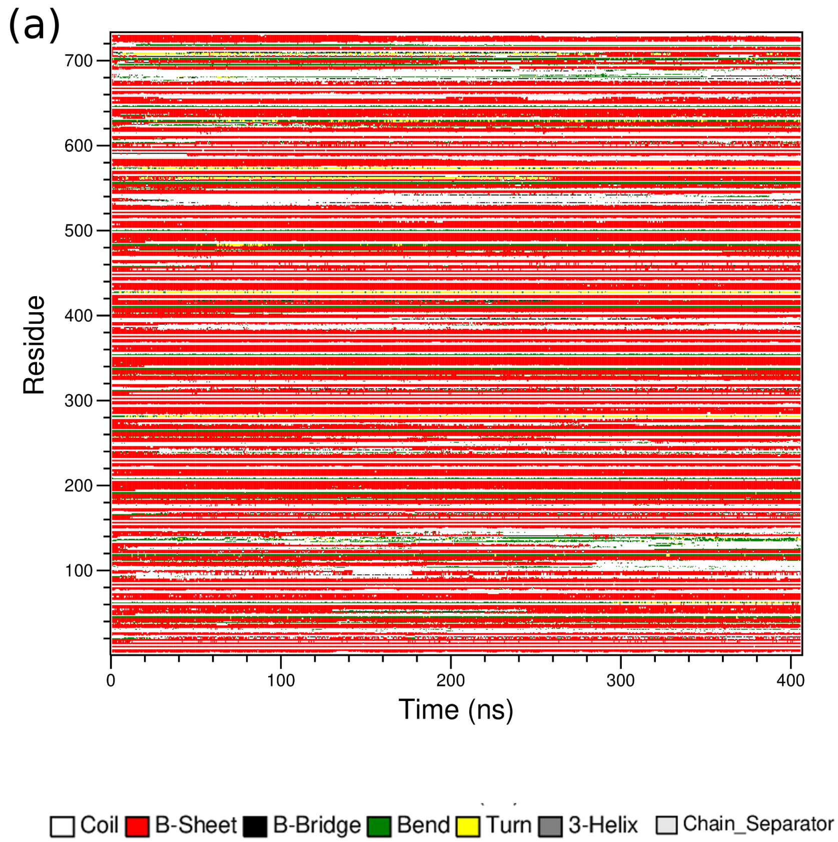


Figure S13: DSSP timelines for the system (a) POPC+POPE.

### 6 Number of $\beta$-sheet residues

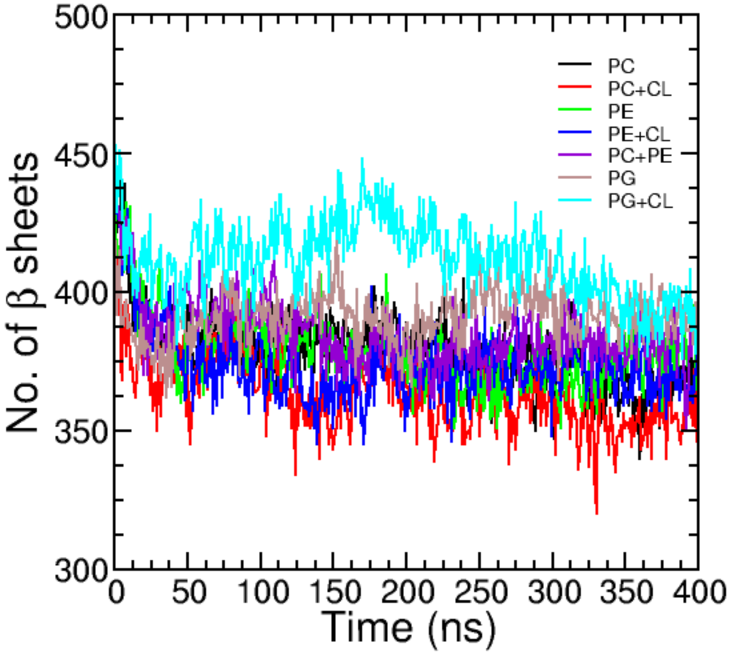


Figure S14: Number of $\beta$-sheet residues for all the systems across the trajectory. PC, PC+CL, PE, PE+CL, PC+PE, PG, PG+CL refers to the POPC, POPC+CHOL, POPE, POPE+CHOL, POPC+POPE, POPG and POPG+CHOL lipid composition respectively.

### 7 Distance Profiles for the CG models

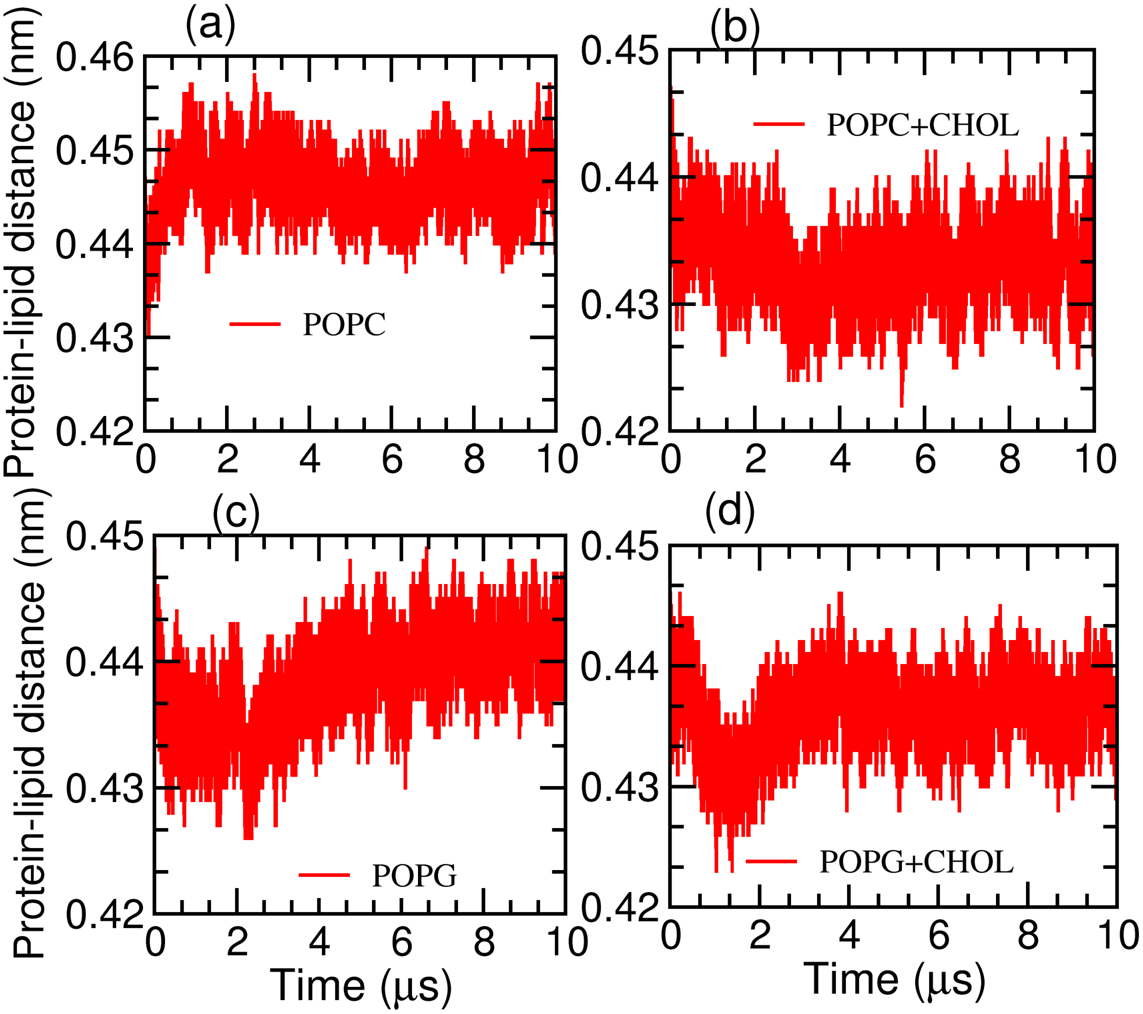


Figure S15: Distance proflies for the center-of-mass between the peptide and lipids (a)-(d).

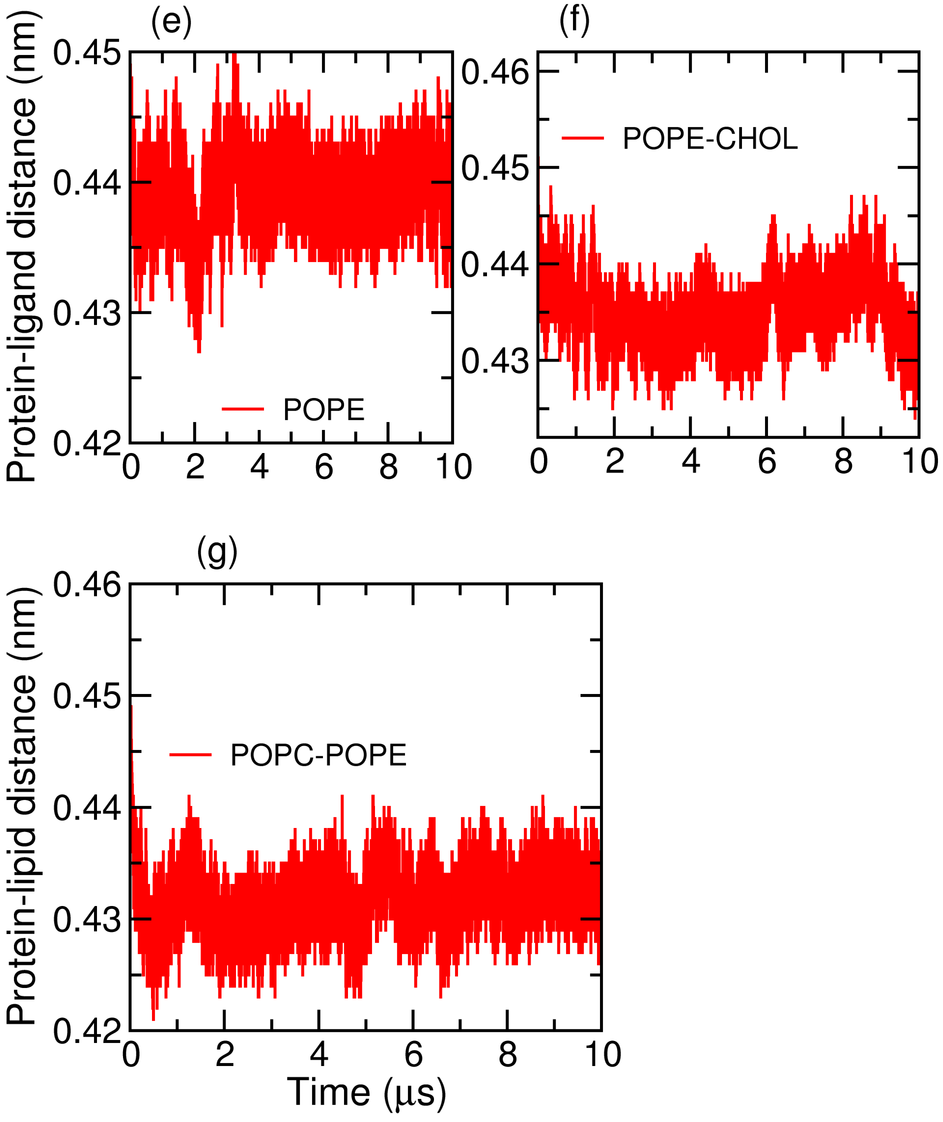


Figure S16: Distance profiles for the center-of-mass between the tau-peptide and lipids (e)-(g).
